# Innate immune control of influenza virus interspecies adaptation

**DOI:** 10.1101/2023.08.23.554491

**Authors:** Parker J. Denz, Samuel Speaks, Adam D. Kenney, Adrian C. Eddy, Jonathan L. Papa, Jack Roettger, Sydney C. Scace, Emily A. Hemann, Adriana Forero, Richard J. Webby, Andrew S. Bowman, Jacob S. Yount

## Abstract

Influenza virus pandemics are caused by viruses from animal reservoirs that adapt to efficiently infect and replicate in human hosts. Here, we investigated whether Interferon-Induced Transmembrane Protein 3 (IFITM3), a host antiviral factor with known human deficiencies, plays a role in interspecies virus infection and adaptation. We found that IFITM3-deficient mice and human cells could be infected with low doses of avian influenza viruses that failed to infect WT counterparts, identifying a new role for IFITM3 in controlling the minimum infectious viral dose threshold. Remarkably, influenza viruses passaged through *Ifitm3*^-/-^ mice exhibited enhanced host adaptation, a result that was distinct from passaging in mice deficient for interferon signaling, which caused virus attenuation. Our data demonstrate that IFITM3 deficiency uniquely facilitates zoonotic influenza virus infections and subsequent adaptation, implicating IFITM3 deficiencies in the human population as a vulnerability for emergence of new pandemic viruses.

## Main

Respiratory virus emergence from animal reservoirs into the human population is a continuous worldwide health threat. The influenza pandemics of 1918, 1957, 1968, and 2009 arose from viruses of avian or swine origin that adapted to replicate robustly in humans^1,2^. Likewise, infection of humans with coronaviruses, such as MERS-CoV, SARS-CoV, and SARS-CoV-2, likely resulted from animal-to-human transmission events followed by virus adaptation to the human respiratory tract and other niches^3^. Despite these notable examples, major outbreaks of new viruses could be considered rare given the routine contact between humans and wildlife throughout the world^4^. This discrepancy suggests that natural impediments to interspecies virus transmission or adaptation exist in human cells. Conversely, genetic defects in such host defense mechanisms could provide opportunities for viruses to enter, and more efficiently adapt to, humans. Here, we tested the hypothesis that deficiencies in the innate immunity protein, Interferon-Induced Transmembrane Protein 3 (IFITM3), facilitate interspecies infection and adaptation by influenza viruses.

Although expressed at low levels in most cell types, IFITM3 is highly upregulated by interferons upon virus infection^5,6^. IFITM3 associates with cell membranes through a transmembrane domain and an *S*-palmitoylated amphipathic helix domain^6,7^. The amphipathic helix alters membrane curvature, lipid composition, and fluidity in a manner that disfavors virus-to-cell membrane fusion, endowing IFITM3 with the ability to block release of virus genomes into the cytoplasm of cells^7-12^. As such, IFITM3 limits infection by a multitude of enveloped viruses *in vitro*, including influenza virus, Zika virus, and SARS-CoV-2^5,13-15^. In humans, there are two deleterious single nucleotide polymorphisms (SNPs) in the *IFITM3* gene that negatively impact its splicing or gene promoter efficacy^16-18^. These SNPs are unexpectedly common in the human population, with 20% of Chinese individuals being homozygous for the splicing SNP (rs12252-C), and 4% of Europeans being homozygous for the promoter SNP (rs34481144-A)^18^. Human IFITM3 deficiencies are associated with heightened severity of influenza, and data are emerging that IFITM3 deficiencies are a risk factor for severe COVID-19^16-20^. Indeed, we and other groups have observed that *Ifitm3*^-/-^ mice experience more severe influenza virus and SARS-CoV-2 infections, mimicking human IFITM3 defects^15,16,21-23^.

Based on these data, and further supported by the recent emergence of new viruses in geographic locations where *IFITM3* SNPs are common (e.g., SARS-CoV, SARS-CoV-2, avian influenza viruses), we posited that IFITM3 deficiency removes a major block to initial zoonotic infection and to subsequent viral replication, thus promoting virus evolution through species-specific adaptive mutations. Herein, we demonstrate that IFITM3 deficiency not only lowers the infectious dose threshold for zoonotic influenza viruses in mice and human cells, but also dramatically promotes influenza virus adaptation to a new species. These functions of IFITM3 could not be inferred from its known roles, as virus passaging in the absence of interferon signaling, which also enhances virus replication, resulted in virus attenuation. Our work indicates that human IFITM3 deficiencies are a unique “Achilles heel” for the emergence of new pandemic viruses and should be considered in pandemic prevention efforts.

### IFITM3 deficiency lowers the minimum infectious dose of avian influenza viruses in mice

Intrinsic immunity is presumed to be among the factors responsible for preventing infection when virus inoculums are below a minimum infectious dose threshold. However, this fundamental concept remains an unproven hypothesis. IFITM3 limits the severity of influenza virus infections in humans and mice^16,17,21-23^, but whether it influences the minimum viral dose required for productive infection *in vivo* has not been tested. We infected WT and *Ifitm3^-/-^* mice with doses of 1, 10, or 50 TCID50 of H5N1 avian influenza virus and measured lung viral loads at day 3 post infection. Infection with 1 TCID50 of H5N1 influenza virus resulted in significant viral replication in the lungs of *Ifitm3^-/-^*mice, approaching titers of 10^4^ TCID50/mL, while live virus was not detected in the lungs of WT mice (**Fig. 1a**). Doses of 10 and 50 TCID50 caused detectable viral loads in both WT and KO mice, with KO mice showing significantly higher viral titers at both doses (**Fig. 1a**). Examination of lung IL-6 as a measure of inflammation induced by viral replication substantiated these titer results. Lung IL-6 was nearly undetectable in WT mice infected with 1 TCID50, but was significantly above baseline in all other samples, including *Ifitm3^-/-^* mice infected with only 1 TCID50 (**Fig. 1b**). We next infected WT and KO mice with a second avian virus using doses of 1 and 10 TCID50. We again observed that 1 TCID50 of H7N3 virus was sufficient to infect *Ifitm3^-/-^* mice as indicated by robust replication and induction of IL-6, whereas the same viral dose did not productively infect WT mice (**Fig. 1c,d**). Dosage of 10 TCID50 resulted in productive virus replication in both WT and *Ifitm3^-/-^*mice, though viral titers and IL-6 induction were each higher in the KO animals (**Fig. 1c,d**).

**Fig. 1.**
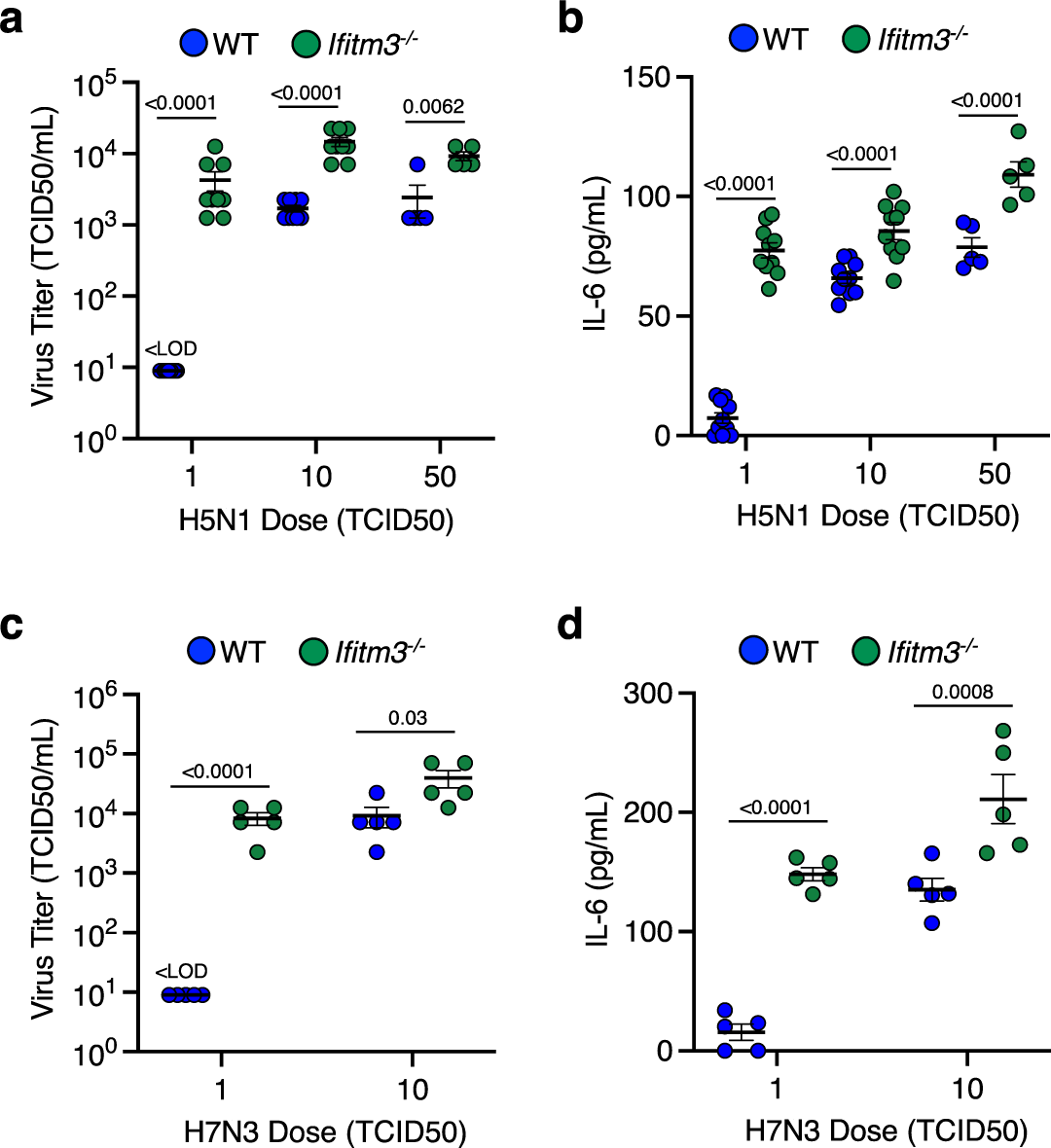
IFITM3 deficiency lowers the minimum infectious dose threshold for avian influenza viruses *in vivo*. WT and *Ifitm3^-/-^* mice were intranasally infected with (**a, b**) 1, 10, or 50 TCID50 of H5N1 avian influenza strain (2 independent experiments for doses 1 and 10 (n=10 mice) and 1 experiment for dose of 50 (n=5 mice)) or with (**c, d**) 1 or 10 TCID50 of H7N3 avian influenza strain (n=5 mice). **a**, **c** Viral titers from lung homogenates at day 3 post infection. **b, d** ELISA quantification of IL-6 levels in lung homogenates at day 3 post infection. **a-d** Error bars represent SEM. Comparisons were analyzed by ANOVA followed by Tukey’s multiple comparisons test. Only comparisons between WT and *Ifitm3^-/-^*mice for each dose are shown.

### IFITM3 deficiency lowers the minimum infectious dose of avian influenza viruses in human cells

To extend our conclusions to a human system, we tested a broad panel of influenza viruses isolated from animals in a relevant human cell type, A549 lung epithelial cells. Specifically, we compared infection rates of 12 avian and swine influenza viruses representing distinct and diverse hemagglutinin protein (HA) subtypes (strain details in **Table S1** and HA-based phylogenetic tree in **Fig. S1.**) in control A549 cells versus those in which IFITM3 was depleted by shRNA knockdown (**Fig. 2a**). The IFITM3-knockdown cells were universally infected at a higher rate (**Fig. 2b,c, Fig. S2**), demonstrating that each of these diverse viruses was restricted by IFITM3 in human cells. Furthermore, IFITM3-knockdown cells maintained higher infection susceptibility than control cells after treatment with type I interferon (**Fig. 2b,c, Fig. S2**), demonstrating that IFITM3 plays a critical and non-redundant role in the human antiviral interferon response against diverse influenza viruses. Since macrophages are also a relevant target of influenza virus^24^, we examined infection of differentiated human THP-1 cells. Once again, infection by the avian and swine viruses was significantly potentiated in the human macrophages lacking IFITM3 with or without interferon treatment (**Fig. 2b,d, Fig. S3**). Increased infection was also observed for IFITM3 KO HAP1 fibroblasts (**Fig. S4**) and for IFITM1/2/3 KO HeLa cells (**Fig. S5**). Of note, though the viruses differed in their sialic acid receptor binding preferences, with several of the viruses showing a preference for α-2,3-linked sialic acid (**Extended Data Table 1**), all the animal-derived viruses were able to infect every cultured human cell type that we tested (**Fig 2b, Fig. S2-S5**). Overall, these data demonstrate that IFITM3 deficiency broadly increases infection of human cells with influenza viruses of animal origin.

**Fig. 2.**
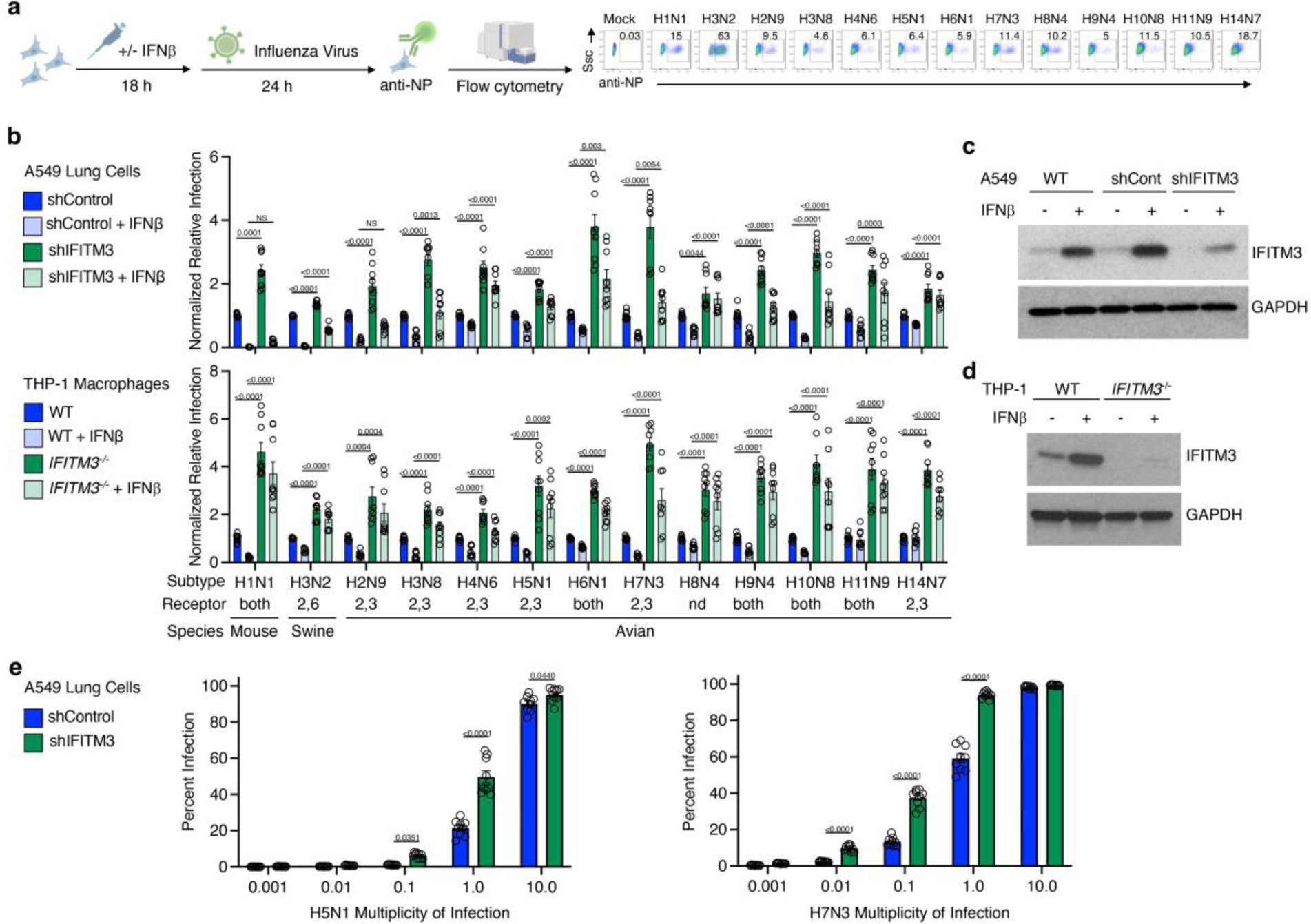
IFITM3 limits zoonotic influenza virus infection of human cells. (**a**) Schematic of *in vitro* infection with potentially zoonotic influenza viruses and representative example from infected A549 human lung cells. (**b**) The indicated A549 cells or THP-1 differentiated macrophages were treated +/- IFNβ for 18 hours, followed by infection with indicated viruses (MOI 1) for 24 hours. Percent infection was determined by flow cytometry and normalized to respective shControl or WT cells without IFNβ pre-treatment. Error bars represent SEM. P values are for the indicated comparisons and were determined by ANOVA followed by Tukey’s multiple comparisons test. Only statistical comparisons between shControl versus shIFITM3 and WT versus *IFITM3^-/-^* are shown. Data are representative of 3 independent experiments each performed in triplicate (n=9). (**c, d**) Western blots of cell lysates at 18 hours +/- IFNβ treatment. Note that commercial IFITM3 antibodies weakly detect IFITM2 in addition to IFITM3. (**e**) The indicated A549 cells were infected with indicated viruses at a range of 0.001 to 10 MOI for 24 hours. Percent infection was determined by flow cytometry. Data are representative of 3 independent experiments each performed in triplicate (n=9). Error bars represent SEM. Comparisons were analyzed by ANOVA followed by Tukey’s multiple comparisons test. Only comparisons between control and knockdown cells are shown at each dose.

To further examine the impact of IFITM3 on minimum infectious dosing we infected A549 lung cells with or without IFITM3 knockdown using H5N1 or H7N3 avian influenza viruses at a range of doses from 0.001 to 10 MOI. We observed that infected cells could be detected at lower viral doses for both H5N1 (**Fig. 2e**) and H7N3 (**Fig. 2f**) when IFITM3 was deficient. Along with data from **Fig. 1**, these data establish that IFITM3 prevents zoonotic influenza virus infection *in vitro* and *in vivo* when the virus dose is below a minimum threshold. These data together demonstrate that IFITM3 increases the minimum infectious dose necessary to achieve a productive influenza virus infection.

### IFITM3 deficiency facilitates adaptation of zoonotic influenza viruses *in vivo*

Several factors have been proposed to favor zoonotic adaptation of a virus to a new host, including levels of virus replication and infectious doses. Given that IFITM3 deficiency impacts both of these features, we tested whether its absence allows influenza virus to adapt more readily *in vivo*. With biosafety in mind, we did not attempt to adapt avian influenza virus to a mammalian host since such adaptations could increase virus replication across mammalian species, including humans, and could thus be considered dual use research of concern^25^. Rather, we sought to adapt viruses of human origin to mice, which has been performed safely for nearly a century in the influenza virus research field^25^. Initially, we performed lung-to-lung passaging of influenza virus strain A/Victoria/361/2011 (H3N2) 10 times through WT or *Ifitm3*^-/-^ mice using intranasal infection and allowing three days for the virus to replicate between successive passages (**Fig. 3a**). We chose this strain because preliminary experiments revealed that it infects mice, but replicates poorly, thereby providing a dynamic range for virus adaptation to a new species.

**Fig. 3.**
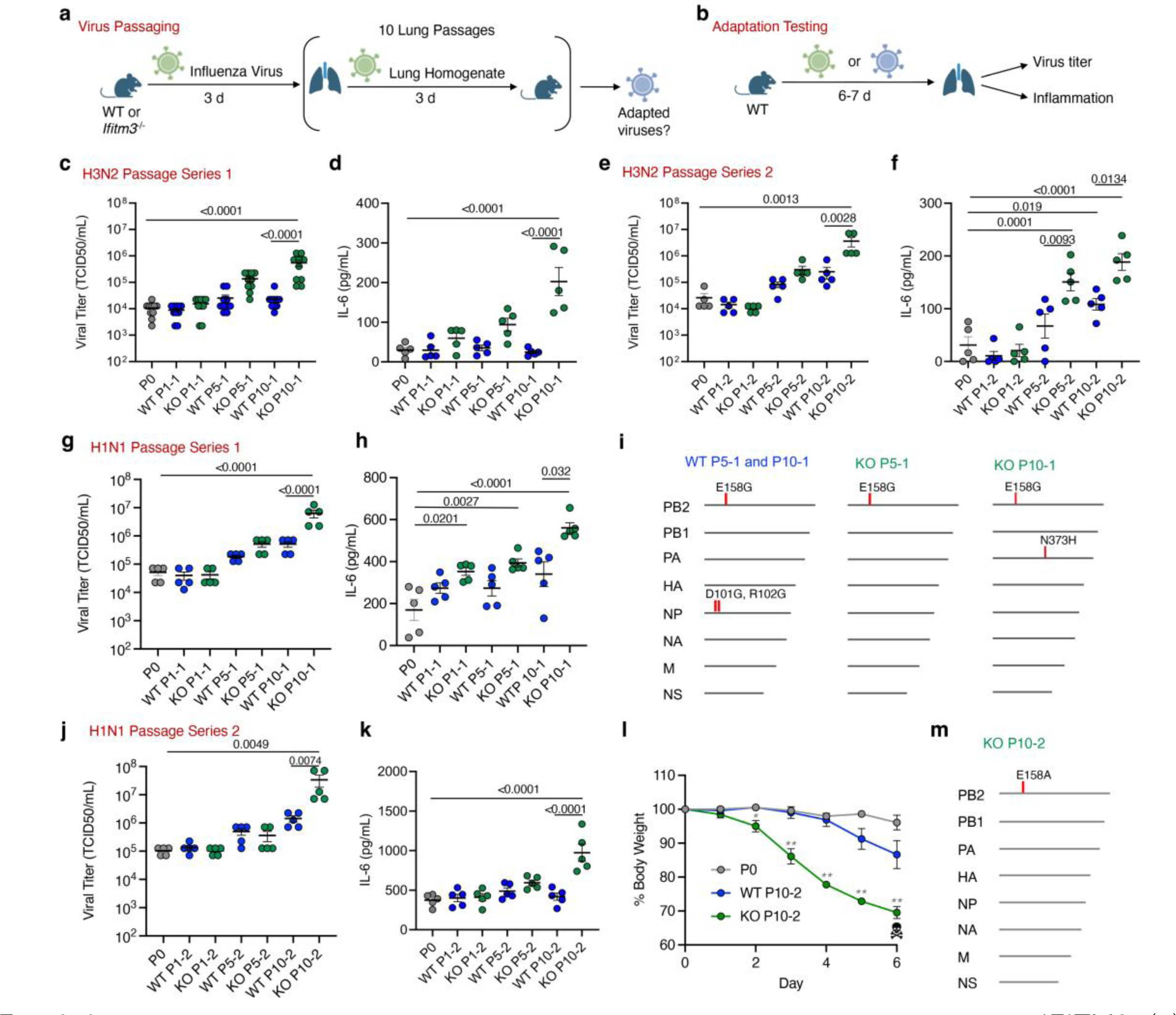
Influenza virus adapts to a new species more readily in the absence of IFITM3. (**a**) Schematic of mouse passaging experiments. Initial intranasal infections were performed with 1,000 TCID50 of parental viruses. (**b**) Schematic of WT mouse challenge with parental or passaged viruses. (**c-h and j-l**) Groups of WT mice were challenged with equal doses of virus passaged 1, 5, or 10 times through WT or *Ifitm3^-/-^*mice and compared to the parent virus (passage 0). (**c, e**) Viral titers from lung homogenates taken at day 7 (**c** represents 2 independent experiments). Error bars represent SD of the mean. Comparisons were analyzed by ANOVA followed by Tukey’s post hoc test. (**d, f**) ELISA quantification of IL-6 levels in lung homogenates of WT and IFITM3 KO mice at day 7 post infection (**d** represents 2 independent experiments). Error bars represent SD of the mean. Comparisons were analyzed by ANOVA followed by Tukey’s multiple comparisons test. (**g, j**) Viral titers from lung homogenates taken at day 7 (**g**) or day 6 (**j**) post infection. Error bars represent SD of the mean. Comparisons were analyzed by ANOVA followed by Tukey’s post hoc test. (**h, k**) ELISA quantification of IL-6 levels in lung homogenates of WT and IFITM3 KO mice at day 7 (**h**) or day 6 (**k**) post infection. Error bars represent SD of the mean. Comparisons were analyzed by ANOVA followed by Tukey’s multiple comparisons test. (**i, m**) Mutations found in the segments of A/California/04/2009 (H1N1) after serial passage through WT or *Ifitm3^-/-^*mice. (**l**) Weight loss. Skull and crossbones indicate humane euthanasia of all animals infected with KO passage 10. Error bars represent SD of the mean, comparisons were made using the Mann-Whitney test (* *P* = 0.0022, ** *P* < 0.0001).

We propagated passages 1, 5, and 10 from WT and KO mice, as well as the parental virus stock (passage 0), in MDCK cells. We then used these expanded stocks to challenge WT mice with equal virus doses (**Fig. 3b**). Lungs from the infected WT animals were collected at day 7 post infection to measure viral titers as a direct indicator of adaptation to mice when compared to the parental virus isolate. We found that, as expected, the ability to replicate in mouse lungs as compared to the parental virus was unchanged in the earliest passages of virus from WT or KO mice (**Fig. 3c**). However, passage 5 from KO mice exhibited an upward trend in viral titers, while KO passage 10 showed a statistically significant, >1 log average increase in lung viral titers (**Fig. 3c**). In contrast, virus passages 5 and 10 from WT mice did not show significantly enhanced replication capacity compared to the parental virus (**Fig. 3c**). As an independent metric for virus adaptation, we measured levels of the inflammatory cytokine IL-6 in lungs during infection, which mirrored viral loads, with IFITM3 KO passage 10 inducing a statistically significant increase in IL-6 compared to infection with parent virus (**Fig. 3d**). Since virus adaptation is a stochastic process involving the random generation and outgrowth of beneficial mutations, we independently repeated our H3N2 virus passaging and testing a second time, and obtained similar results indicating enhanced adaptation occurring in IFITM3 KOs (**Fig. 3e,f**). Overall, our data from both passage series demonstrate that influenza virus reproducibly adapted more readily and efficiently, i.e., gained enhanced replication capacity and induction of inflammation, when passaged through IFITM3 deficient versus WT hosts.

We next tested whether IFITM3 deficiency is unique in its ability to facilitate virus adaptation or whether passaging in another immune compromised system that allows enhanced virus replication would also promote interspecies adaptation. The interferon system is particularly important in this regard as interferons widely inhibit virus replication^26^. Further, it has become increasingly apparent throughout the recent COVID-19 pandemic that genetic and antibody-mediated interferon deficiencies are significantly more common in the human population than previously appreciated^27,28^. We thus repeated our passaging and challenge regimen with influenza virus A/Victoria/361/2011 (H3N2) comparing passaging through WT versus *Stat1^-/-^*mice, which lack signaling downstream of all interferon receptors (**Fig. S6a,b**). Consistent with our previous results, we observed no significant adaptation of virus passaged through WT mice (**Fig. S6c**). In contrast, passage 5 from the STAT1 deficient mice exhibited a downward trend in viral replication, while *Stat1^-/-^* passage 10 showed a statistically significant reduction in the ability of passaged virus to replication in WT mice (**Fig. S6c**). The titer data were supported by measurements of inflammatory cytokine IL-6, in that STAT1 KO passage 10 showed a significant decrease in cytokine induction compared to other groups (**Fig. S6d**). Our findings are consistent with the principle that, in the absence of selective pressures from the interferon system, viral interferon antagonism mechanisms become inactive through mutation, resulting in virus attenuation in WT systems. Importantly, these results underscore how IFITM3 may be a unique vulnerability in innate immune response pathways that, when impaired, can promote viral adaptation to new hosts rather than viral attenuation.

To further investigate the generality of our findings on IFITM3, we repeated the passaging regimen in WT and *Ifitm3*^-/-^ mice using pandemic influenza virus strain A/California/04/2009 (H1N1). Virus from passages 5 or 10 in WT mice were modestly enhanced in replicative capacity (**Fig. 3g**) without significant changes in their IL-6 induction capacity (**Fig. 3h**). In these experiments, we sequenced viruses and found that the enhanced replication of the WT passages was associated with two amino acid changes in the viral NP (D101G, R102G) and a single change in PB2 (E158G) that is a known polymerase activity-increasing adaptive mutation selected in mice^29,30^. Consistent with our previous results, virus passaged in *Ifitm3*^-/-^ mice replicated to higher titers compared to the parental strain and to the respective WT mouse passages (**Fig. 3g**). In addition to the highest viral titer, KO passage 10 also induced robust levels of IL-6 in infected lungs (**Fig. 3h**). KO passages 5 and 10 possessed the same adaptive PB2 E158G mutation seen in WT passages 5 and 10 (**Fig. 3i**). However, the heightened replication and induction of IL-6 by KO passage 10 relative to KO passage 5 was associated with a single change in the PA polymerase subunit (N373H), representing a novel mutation not previously reported in mouse adaptation studies (**Fig. 3i**). In a second series of passages with the H1N1 virus, the most robust adaptation in terms of viral replication and IL-6 induction was again seen for KO passage 10 (**Fig. 3j,k**). Remarkably, this viral stock became highly virulent, with all infected mice meeting humane endpoint criteria of 30% weight loss by day 6 post infection (**Fig. 3l**). The virulence of this KO passage 10 was linked to a single genetic alteration, specifically an E158A mutation in PB2 (**Fig. 3m**). This mutation was previously found to increase viral polymerase activity above that caused by the PB2 E158G detected in our first series of H1N1 passages^30^. These results highlight *Ifitm3^-/-^* mice as a tool to rapidly adapt influenza viruses to mice and to explore adaptive mutation strategies. Together with data for the H3N2 virus, our H1N1 passaging experiments demonstrate that adaptation of influenza viruses to a new host species is accelerated in the absence of IFITM3, fueled by mutations that readily emerge in the compromised host.

## Discussion

Three fundamentally important discoveries about IFITM3 emerged from our study. First, IFITM3 raises the minimal infectious dose of influenza virus required to achieve a detectable infection in human cells, as well as a productive infection *in vivo*. Second, IFITM3 serves as a critical barrier to pandemic virus emergence by limiting adaptation of influenza viruses to new host species, whereas global defects in interferon responses lead to virus attenuation. Third, IFITM3, whether basally expressed or induced by interferon, broadly restricts infection of human cells by potentially zoonotic influenza viruses, and this function was independent of viral receptor binding preferences. Each of these findings support the premise that human IFITM3 deficiencies represent a critical vulnerability for the entry and adaptation of zoonotic influenza viruses in the population.

Our experiments also indirectly address transmission of influenza viruses. Mice are unable to transmit influenza virus to one another, even when in close contact^25^. We have also observed in unpublished work that IFITM3 KO mice, perhaps the most susceptible mouse model of influenza, do not transmit mouse adapted influenza virus or mouse adapted SARS-CoV-2 to non-infected, cohoused KO animals. Nonetheless, we found that the minimal *in vivo* infectious dose of avian influenza virus, a critical aspect of the transmission process, was lowered in the absence of IFITM3. Specifically, *Ifitm3*^-/-^ animals showed significant replication of virus in their lungs after infection with only 1 TCID50 of H5N1 or H7N3 virus, while titers were undetectable in WT lungs when infected at the same dose. These results provide hard data for the long-presumed concept that minimum infectious doses of a pathogenic virus are set by levels of resistance from the innate immune system. Specifically, we show that IFITM3 underlies sterilizing immunity that prevents infection at low influenza virus doses that are below a minimum threshold.

We designed our virus passaging experiments to model transmission of a zoonotic virus through a family or community in which deleterious *IFITM3* SNPs are prevalent. One caveat to this approach is that, while our engineered mice are fully devoid of IFITM3 production, deleterious SNPs in human *IFITM3* substantially reduce IFITM3 levels or activity, but do not ablate the protein^16,17^. In this regard, lung cells partially deficient in IFITM3 (IFITM3-knockdown A549 cells in **Fig. 2**) are significantly more susceptible to diverse influenza virus strains. Thus, while infections in humans with IFITM3 deficiencies may represent more nuanced scenarios, experimentation with complete KO mice allowed us to unambiguously identify important and relevant functionality of IFITM3 in limiting interspecies infection and adaptation of zoonotic viruses *in vivo*, a key finding that is likely germane to preventing pandemics in humans.

RNA viruses generally adapt to new host species by producing mutations arising from error-prone polymerases ^31^. Beneficial mutations that allow the virus to replicate more efficiently are enriched by outcompeting the parental virus and other mutants. Indeed, influenza virus and several other viruses replicate to higher titers in *Ifitm3*^-/-^ versus WT mice^15,16,21-23,32-34^, which represents a plausible mechanism for enhanced adaptation in these animals^31^. In our current study, mutations in the 2009 pandemic H1N1 virus passages that drive adaptation in mice were primarily observed in the viral polymerase subunits. Indeed, PB2 mutations at the E158 position are consistently reported in mouse-adapted versions of this virus^29,30^, with the E158A mutation leading to the most dramatic enhancement of polymerase activity^30^. Moreover, the relative impact of E158 mutations on polymerase activity are in accord with viral replication and virulence levels observed in our *in vivo* experiments. We also identified a novel N373H mutation in PA from one H1N1 passage series, which emerged subsequent to the E158G mutation. The dual mutations were associated with enhanced replication compared to E158G alone. The mechanistic impact of the PA N373H mutation remains to be determined and may provide future insights into influenza virus replication and adaptation.

The prevalence of genetic and immunological mechanisms that impair interferon induction or signaling in humans has garnered increased attention during the COVID-19 pandemic, since 20% of all severe COVID-19 cases could be attributed to interferon defects ^27,28^. Given that IFITM3 expression is induced by interferon, such defects likely diminish its induction, as well as that of other antiviral proteins, which could allow increased replication of viruses and enhanced adaptation to humans. However, passaging through interferon deficient systems often results in attenuated viruses that have lost interferon antagonism due to a lack of selective pressure for their maintenance^35,36^. Indeed, our passaging of influenza virus in *Stat1*^-/-^ animals resulted in virus attenuation, despite influenza virus replicating to high titers in interferon deficient systems^37^. Additional studies on viral evolution in human populations and engineered animals will be required to address how broad interferon defects mechanistically influence adaptive evolution of influenza virus and other pathogens. To our knowledge, influenza viruses capable of fully evading IFITM3 restriction have not been identified to date. Likewise, unlike the well characterized antagonism of interferon by the influenza virus NS1 protein^37^, viral antagonism of IFITM3 has not been reported. Thus, in contrast to interferon, IFITM3 is unlikely to contribute to selective pressures on influenza virus, allowing the virus to efficiently adapt in its absence. IFITM3 deficiency may therefore represent a particularly unique vulnerability for the entry and adaptation of viruses in a new host species. Broad testing of IFITM3 status in the human population could bolster pandemic prevention efforts by allowing vulnerable individuals to receive greater vaccine coverage or exercise enhanced precautions when encountering animal reservoirs of infection.

## Materials and Methods

### Cell culture

Control and IFITM3 knockdown A549 cells were generated by lentiviral shRNA-mediated targeting using previously described constructs and methods^38^. THP-1 IFITM3 knockout cells were generated via CRISPR-Cas9 targeting (provided by Dr. Anasuya Sarkar of the Ohio State University). HeLa IFITM1/2/3 knockout and HAP1 IFITM3 knockout were purchased from ATCC (CRL-3452) and Horizon Discovery Biosciences (HZGHC004186c010), respectively. MDCK cells were obtained from BEI Resources (NR-2628). Cells were maintained in DMEM medium (Fisher scientific) supplemented with 10% EquaFETAL bovine serum (Atlas Biologicals) except for the THP-1 cells which were maintained in RPMI 1640 medium (Fisher scientific) supplemented with 10% EquaFETAL bovine serum (Atlas Biologicals). All cells were cultured in a humidified incubator at 37 °C with 5% CO2.

### Virus stocks and in vitro infection

Influenza viruses A/Puerto Rico/8/34 (H1N1) (PR8, provided by Dr. Thomas Moran of the Icahn School of Medicine at Mt. Sinai), A/Victoria/361/2011 (H3N2) (BEI Resources), and A/California/4/2009 (H1N1) (BEI Resources) were propagated in 10-day embryonated chicken eggs (Charles River Laboratories) and titered on MDCK cells. The potential zoonotic viruses were collected by Andrew Bowman of the Ohio State University and were also propagated in embryonated chicken eggs (Charles River Laboratories) and titered on MDCK cells. Infection of cell lines was done at an MOI of 1 for 24hrs. Cells that received Interferon-β pre-treatment were stimulated for 18 hours with 100 units of recombinant human Interferon-β protein (Millipore, IF014) prior to infection.

### Mouse studies

All mice used in this study were of the C57BL/6J background. *Ifitm3^-/-^* mice with a 53 base pair deletion in exon 1 of the *Ifitm3* gene were described previously^23^. *Stat1^-/-^* mice were obtained from The Jackson Laboratory (Strain #: 012606). All animals were housed in sterile ventilated cages within a specific pathogen-free environment. Male and female mice between 6 and 10 weeks of age were used in our experiments. All mouse infections were performed intranasally under anesthesia with isoflurane. For virus passaging, mice were initially infected with doses of 1,000 TCID50, and mouse organs were collected and homogenized in 500μL of sterile saline. Propagation and titering of passaged viruses, as well as parental control viruses, was done on MDCK cells. WT mice were infected with equal doses (1,000 TCID50) for testing of virus adaptation. For determining lung titers and cytokine levels in virus adaptation studies, lungs were collected and homogenized in 1ml of PBS, flash-frozen, and stored at -80°C prior to titering on MDCK cells or analysis via ELISA. The generation of mouse-adapted viruses via *in vivo* passaging was evaluated and approved by the OSU Institutional Biosafety Committee (protocol # 2016R00000014-R1). All infected mice and influenza virus-containing samples were handled with Biosafety Level 2 precautions, including the use of biosafety cabinets and personal protective equipment. Laboratory staff members were also vaccinated annually against influenza A and B viruses. Mouse procedures were approved by the OSU Institutional Animal Care and Use Committee (protocol # 2016A00000051-R2) and were performed in accordance with guidelines for the ethical use of animals.

### Virus Sequencing

Viral RNA was extracted from MDCK-grown stocks using the RNeasy Mini Kit (Qiagen), and cDNA was synthesized using the Superscript IV First-Strand Synthesis System (Invitrogen). The influenza A virus gene segments were amplified using modified universal primers in a multi-segment PCR as previously described^39^. PCR products were purified using Agencourt AMPure XP beads according to the manufacturer’s protocol (BeckmanCoulter). Libraries were prepared using the Nextera XT DNA Library Prep Kit (Illumina) according to the manufacturer’s protocol and sequenced using a MiSeq Reagent Kit v2 (300 cycles) on a MiSeq System (Illumina). Sequencing reads were then quality trimmed and assembled using CLC Genomics Workbench (version 22.0.1).

### Western blotting

For detection of protein expression in cell lines, samples were lysed in 1% SDS buffer (0.1 mM triethanolamine, 150 mM NaCl, 1% SDS (Sigma), pH 7.4) containing Roche Complete Protease Inhibitor Cocktail. The lysates were centrifuged at 15,000 rpm for 10 min and soluble protein supernatants were used for Western blot analysis. Equal amounts of protein (30ug) were separated by SDS-PAGE and transferred onto membranes. Membranes were blocked with 10% non-fat milk in Phosphate-buffered saline with 0.1% Tween-20 (PBST) and probed with antibodies against IFITM1 (Cell Signaling Technology, #13126), IFITM2 (Cell Signaling Technology, #13530), IFITM3 (ProteinTech, 11714-1-AP), and GAPDH (Thermo Scientific, ZG003).

### IL-6 ELISAs

IL-6 ELISAs were performed on supernatant from mouse lung homogenates using the Mouse IL-6 Duoset ELISA kit from R&D Systems (catalog # DY406) according to manufacturer’s instructions.

### Flow cytometry

Cells were fixed with 4%paraformaldehyde for 10 minutes at room temperature, permeabilized with 0.1% Triton X-100 in PBS, stained with an antibody against Influenza A Virus Nucleoprotein (Abcam, ab20343) conjugated with Goat anti-Mouse IgG (H+L) Highly Cross-Adsorbed Secondary Antibody, Alexa Fluor™ 647 (Thermo Scientific, Catalog # A-21236) resuspended in 2% FBS in PBS, and run on a BD FACS Canto II flow cytometer. Results were analyzed using FlowJo Software (version 10.8.1).

## Author Contributions

P.J.D. organized and performed all experiments in the manuscript, analyzed data, and contributed to manuscript writing and editing of the manuscript. S.S., A.D.K., A.C.E., J.L.P., J.R., and S.S. assisted with mouse colony genotyping/maintenance, virus titering, and mouse experiments. E.A.H. and A.F. provided conceptual input and experimental design and edited the manuscript. R.J.W. performed 2009 H1N1 virus sequencing and analysis. A.S.B. provided zoonotic influenza virus strains and conceptual input. J.S.Y. conceived the study, supervised experimental design and performance of experiments, analyzed and interpreted data, and wrote the manuscript.

## Acknowledgments

The authors thank Dr. Eugene Oltz (The Ohio State University) for critical reading and editing of the manuscript. We thank Dr. Ana Sarkar and Paul Consiglio (The Ohio State University) for providing IFITM3 KO THP-1 cells. Research in the Yount Laboratory is funded by NIH grants AI130110, HL168501, HL157215, HL154001, and AI151230. Research in the Hemann laboratory is funded by NIH grant AI146141. A.D.K. and P.J.D. were funded by NIH training grant AI112542.

## Data availability

All data are available in the main text or the supplementary materials.

---

**Fig. S1.**
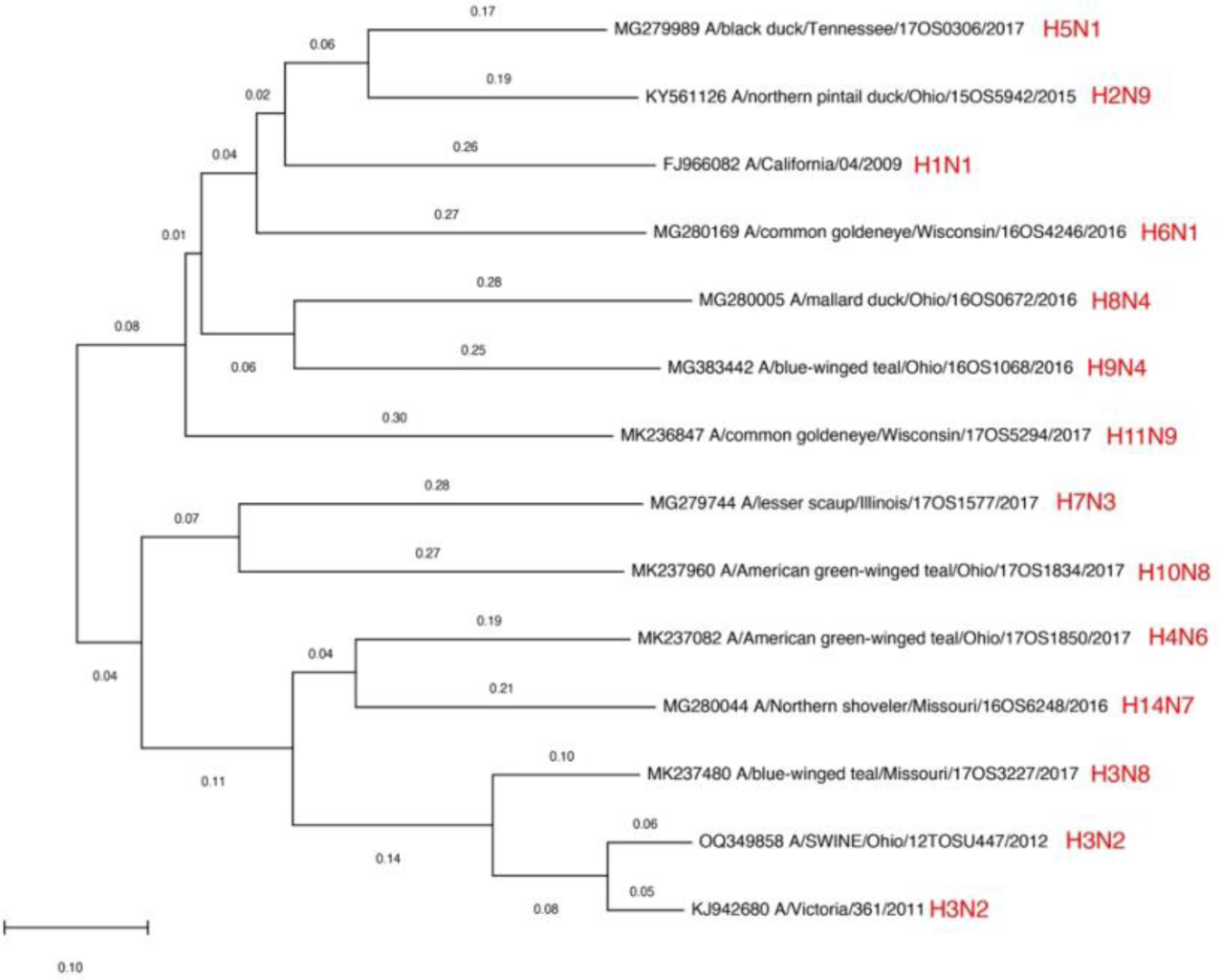
Phylogenetic tree of avian-, swine-, and human-derived influenza viruses utilized in this study. Alignments and tree estimates for whole HA genomic segments were performed using Molecular Evolutionary Genetics Analysis (MEGA) software. Numbers represent the number of nucleotide substitutions normalized to the length of the sequences.

**Fig. S2.**
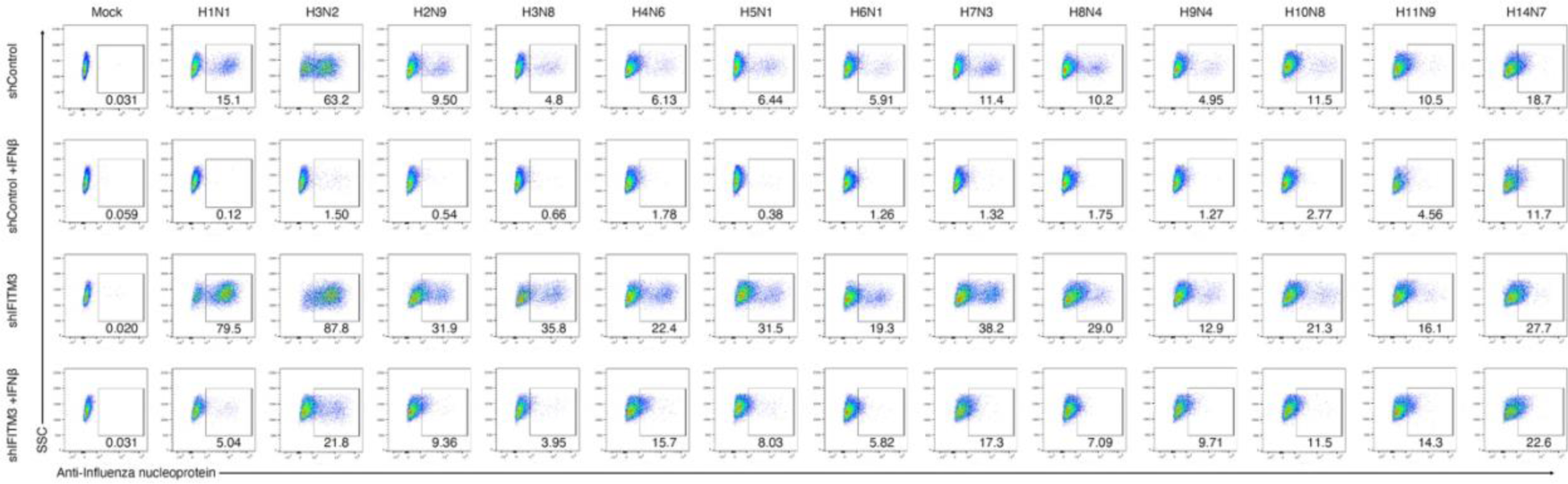
Example raw flow cytometry data for determination of normalized infection of control (shControl) and IFITM3-knockdown (shIFITM3) A549 lung cells +/- IFNb treatment with each of the indicated virus strains as in main text Figure 1. SSC, side scatter; Anti-influenza nucleoprotein antibody was detected in the APC channel with secondary antibody labeled with Alexafluor 647.

**Fig. S3.**
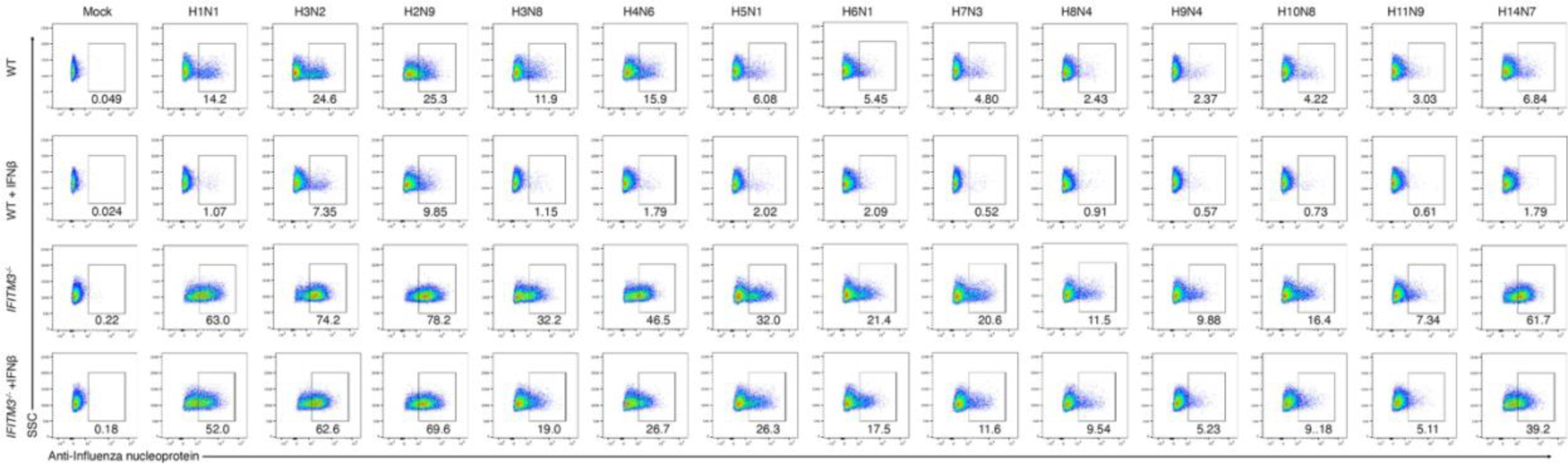
Example raw flow cytometry data for determination of normalized infection of WT and *IFITM3^-/-^*THP-1 macrophages +/- IFNb treatment with each of the indicated virus strains as in main text Figure 1. SSC, side scatter; Anti-influenza nucleoprotein antibody was detected in the APC channel with secondary antibody labeled with Alexafluor 647.

**Fig. S4.**
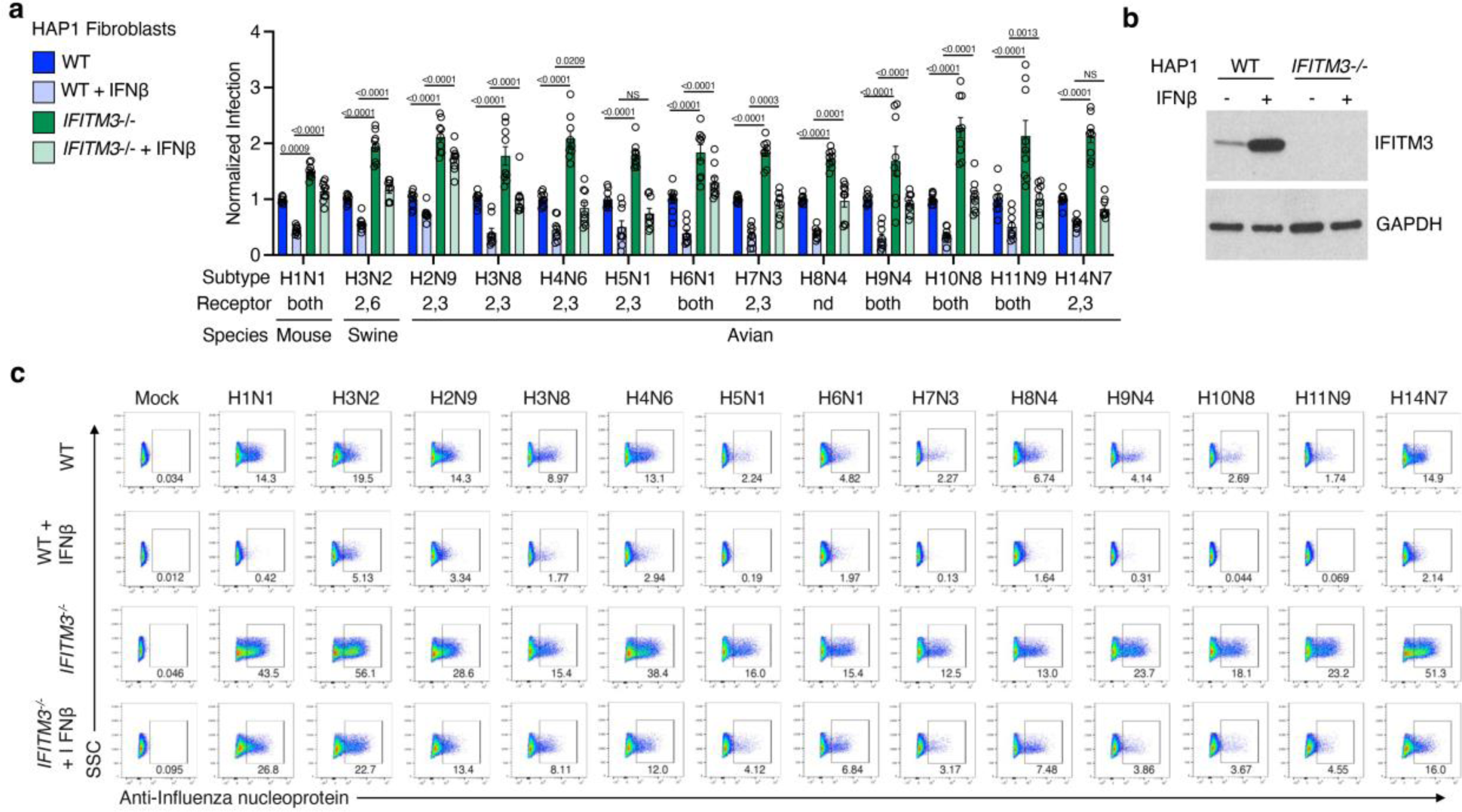
IFITM3 limits zoonotic influenza virus infection of human fibroblast cells. HAP1 cells were treated +/- IFNβ for 18 hours, followed by infection with indicated viruses (MOI 1) for 24 hours. (**a**) Percent infection was determined by flow cytometry and normalized to results for WT cells without IFNβ pre-treatment. Error bars represent SEM. P values are for the indicated comparisons and were determined by ANOVA followed by Tukey’s multiple comparisons test. Only statistical comparisons between WT versus *IFITM3^-/-^*are shown. Data are representative of 3 independent experiments each performed in triplicate (n=9). (**b**) Western blots of cell lysates at 18 hours +/- IFNβ treatment. (**c**) Example raw flow cytometry data for determination of normalized infection of WT and *IFITM3^-/-^*HAP1 cells +/- IFNb treatment with each of the indicated virus strains as in (**a**). SSC, side scatter; Anti-influenza nucleoprotein antibody was detected in the APC channel with secondary antibody labeled with Alexafluor 647.

**Fig. S5.**
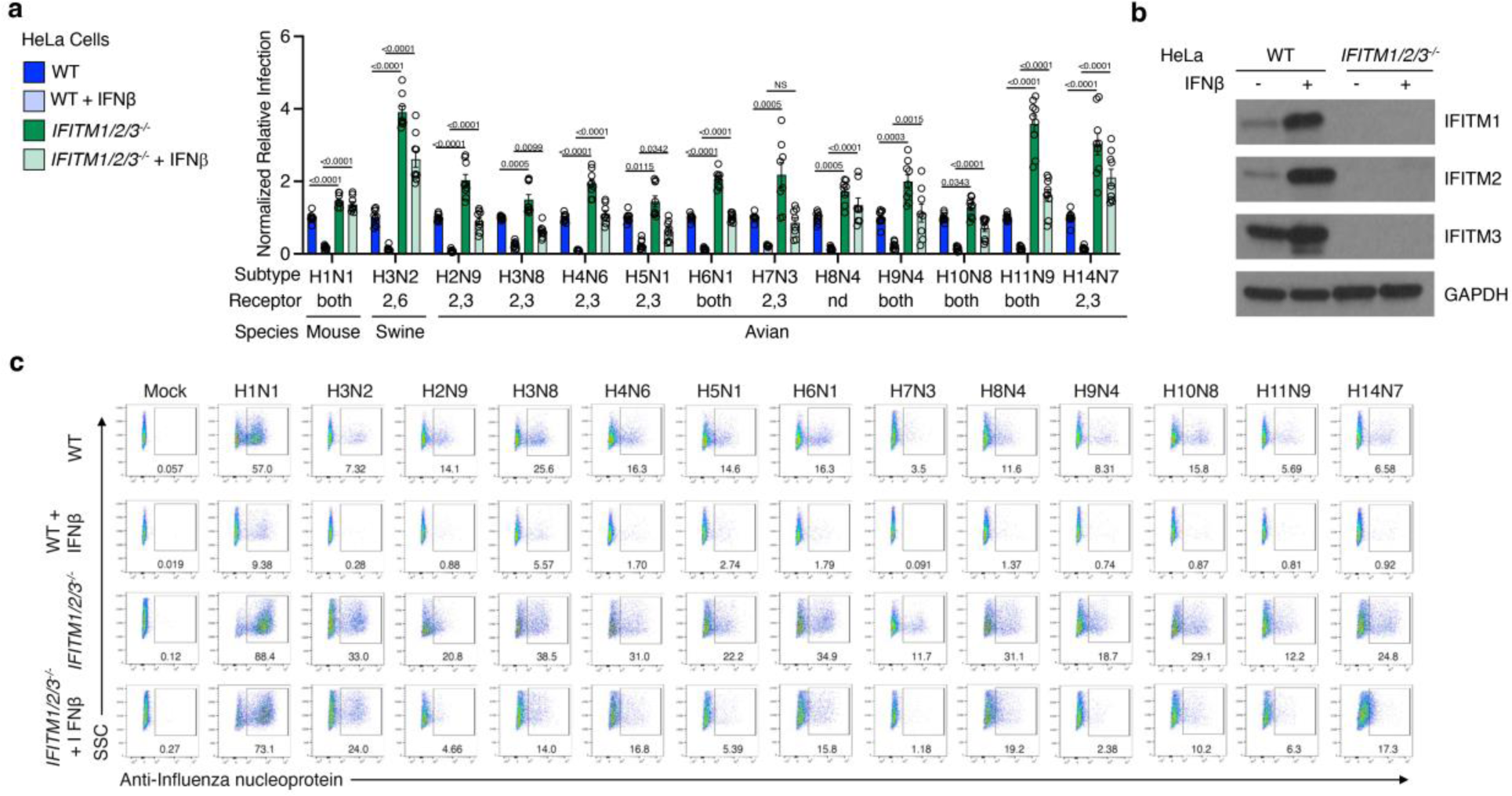
IFITM3 limits zoonotic influenza virus infection of HeLa cells. HeLa cells were treated +/- IFNβ for 18 hours, followed by infection with indicated viruses (MOI 1) for 24 hours. (**a**) Percent infection was determined by flow cytometry and normalized to results for WT cells without IFNβ pre-treatment. Error bars represent SEM. P values are for the indicated comparisons and were determined by ANOVA followed by Tukey’s multiple comparisons test. Only statistical comparisons between WT versus *IFITM3^-/-^*are shown. Data are representative of 3 independent experiments each performed in triplicate (n=9). (**b**) Western blots of cell lysates at 18 hours +/- IFNβ treatment. (**c**) Example raw flow cytometry data for determination of normalized infection of WT and *IFITM1/2/3^-/-^* HeLa cells +/- IFNb treatment with each of the indicated virus strains as in **a**. SSC, side scatter; Anti-influenza nucleoprotein antibody was detected in the APC channel with secondary antibody labeled with Alexafluor 647.

**Fig. S6.**
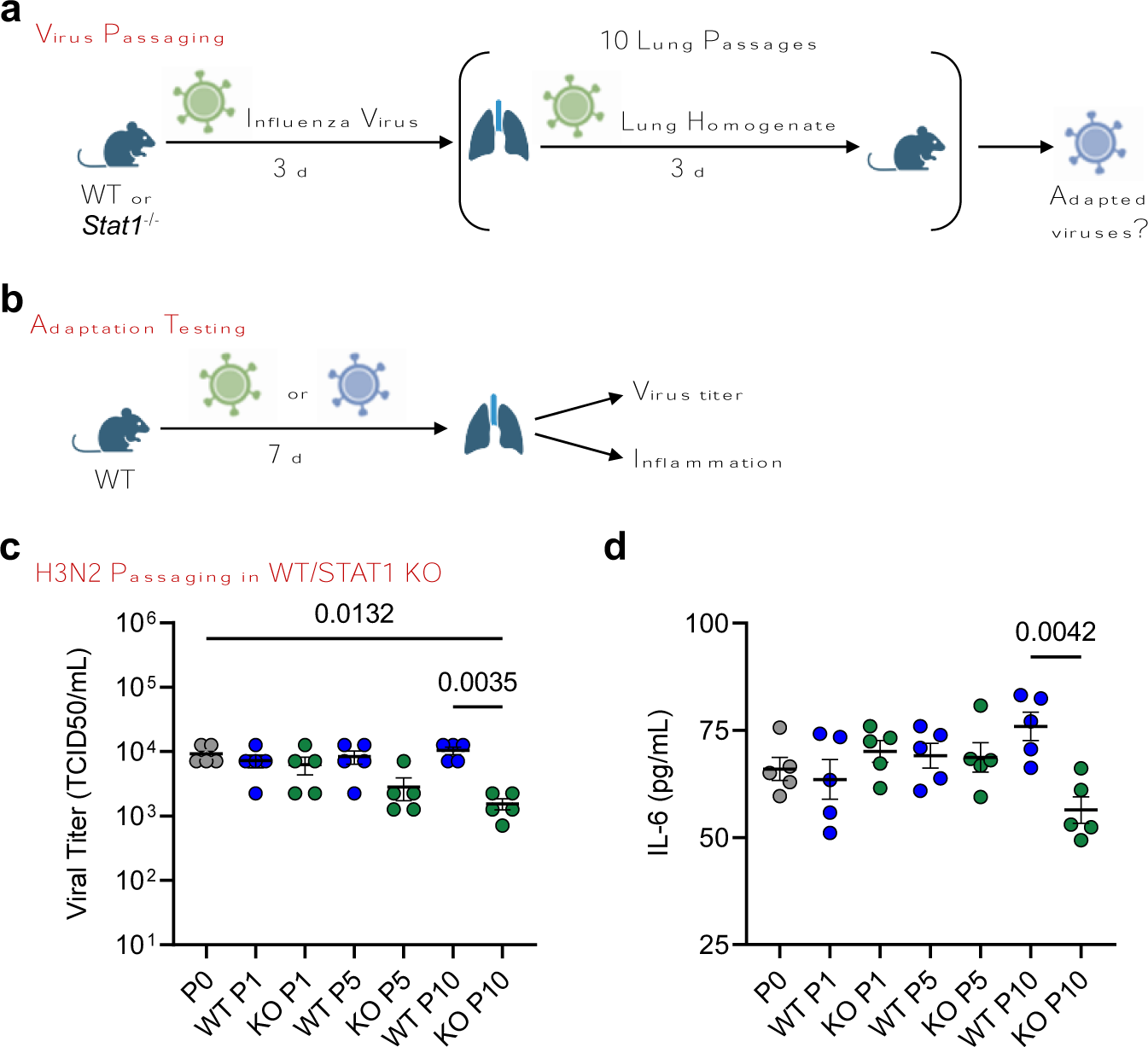
Influenza virus is attenuated when passaged in the absence of STAT1. (**a**) Schematic of mouse passaging experiments. Initial intranasal infections were performed with 1,000 TCID50 of parental viruses. (**b**) Schematic of WT mouse challenge with parental or passaged viruses. (**c,d**) Groups of WT mice were challenged with equal doses of virus passaged 1, 5, or 10 times through WT or *Stat1^-/-^*mice and compared to the parent virus (passage 0). (**c**) Viral titers from lung homogenates taken at day 7 post infection. Error bars represent SEM. Comparisons were analyzed by ANOVA followed by Tukey’s multiple comparisons test. (**d**) ELISA quantification of IL-6 levels in lung homogenates of WT and IFITM3 KO mice at day 7 post infection. Error bars represent SEM. Comparisons were analyzed by ANOVA followed by Tukey’s multiple comparisons test.

**Table S1.** Virus strains utilized in this study and their sialic acid receptor binding preferences. Full strain information and sample identifiers are listed. Data shown are hemagglutination titers used to determine virus sialic acid receptor binding preferences. TRBC, turkey red blood cell; Treated TRBC indicates TRBC incubated with enzyme to remove a-2,3 sialic acid linkages; HRBC, human red blood cell; NT, not tested; ND, not determinable.

| Influenza Virus Strain | Sample ID | Treated TRBC (α-2,6) Titer | TRBC (α-2,6/α-2,3) Titer | HRBC (α-2,3) Titer | Receptor Preference |
| --- | --- | --- | --- | --- | --- |
| A/Puerto Rico/8/1934 (H1N1) mouse adapted | - | NT | NT | NT | Both* |
| A/Swine/Ohio/2012 (H3N2) | 12TOSU0447 | 4069 | 1024 | 8 | α-2,6 |
| A/northern pintail duck/Ohio/2015 (H2N9) | 15OS5942 | <2 | 64 | 32 | α-2,3 |
| A/blue-winged teal/Missouri/2017 (H3N8) | 17OS3227 | <2 | 512 | 32 | α-2,3 |
| A/American green-winged teal/Ohio/2017 (H4N6) | 17OS1850 | <2 | 1024 | 1024 | α-2,3 |
| A/black duck/Tennessee/2017 (H5N1) | 17OS0306 | <2 | 32 | 32 | α-2,3 |
| A/common goldeneye/Wisconsin/2016 (H6N1) | 16OS4246 | 32 | 32 | 32 | Both |
| A/lesser scaup/Illinois/2017 (H7N3) | 17OS1577 | 4 | 128 | 128 | α-2,3 |
| A/mallard duck/Ohio/2016 (H8N4) | 16OS0672 | <2 | <2 | <2 | ND |
| A/blue-winged teal/Ohio/2016 (H9N4) | 16OS1068 | 64 | 64 | 32 | Both |
| A/American green-winged teal/Ohio/2017 (H10N8) | 17OS1834 | 8 | 8 | 8 | Both |
| A/common goldeneye/Wisconsin/2017 (H11N9) | 17OS5294 | 2048 | 2048 | 2048 | Both |
| A/northern shoveler/Missouri/2016 (H14N7) | 16OS6248 | <2 | 32 | 32 | α-2,3 |

\*Receptor preference for the A/Puerto Rico/8/1934 (H1N1) virus of the Mt. Sinai lineage is from Meng, et al., *Influenza and Other Respiratory Viruses*, 2010.

